# The evolution of cold nociception in drosophilid larvae and identification of a neural basis for cold acclimation

**DOI:** 10.1101/2021.01.04.425280

**Authors:** Nathaniel J. Himmel, Jamin M. Letcher, Akira Sakurai, Thomas R. Gray, Maggie N. Benson, Kevin J. Donaldson, Daniel N. Cox

## Abstract

Cold temperatures can be fatal to insects, but many species have evolved the ability to cold acclimate, thereby increasing their cold tolerance. While there is a growing body of knowledge concerning the mechanisms underlying cold tolerance, relatively little is known concerning how insects sense noxious cold (cold nociception), or how cold nociception might function in cold tolerance. It has been previously shown that *Drosophila melanogaster* larvae perform highly stereotyped, cold-evoked behaviors under the control of noxious cold-sensing neurons (nociceptors) innervating the barrier epidermis. In the present study, we first sought to describe cold-nociceptive behavior among 11 drosophilid species with differing cold tolerances and from differing climates. Behavioral analyses revealed that the predominant cold-evoked response among drosophilid larvae is a head-to-tail contraction (CT) behavior, which is likely inherited from a common ancestor. However, despite lack of phylogenetic signal (suggesting trait lability), the CT behavior was transient and there was no clear evidence that cold sensitivity was related to thermal environment; collectively this suggests that the behavior might not be adaptive. We therefore sought to uncover an alternative way that cold nociception might be protective. Using a combination of cold-shock assays, optogenetics, electrophysiology, and methods to genetically disrupt neural transmission, we demonstrate that cold sensing neurons in *Drosophila melanogaster* (Class III nociceptors) are sensitized by and critical to cold acclimation. Moreover, we demonstrate that cold acclimation can be optogenetically-evoked, *sans* cold. Collectively, these findings reveal that cold nociception constitutes a peripheral neural basis for *Drosophila* larval cold acclimation.

**Significance Statement:** Many insects adapt to cold in response to developmental exposure to cool temperatures. While there is a growing body of knowledge concerning the mechanisms underlying cold tolerance, it is unknown how sensory neurons might contribute. Here, we show that noxious cold sensing (cold nociception) is widely present among drosophilid larvae, and that cold-sensing neurons (Class III cold nociceptors) are necessary and sufficient drivers of cold acclimation. This suggests that cold acclimation has, at least in part, a neural basis.

## Introduction

Small ectotherms can be particularly susceptible to damage by chilling. For example, in *Drosophila melanogaster*, prolonged exposure to extreme cold can cause permanent defects in fertility, and even death, particularly among early life stages (1, 2). In response, many ectotherms have evolved the ability to adapt to cold, both rapidly (rapid cold hardening; RCH) and in response to long-term, often seasonal, cooling (cold acclimation) (3, 4). Understanding these processes is important to understanding ectotherm ecology and evolution, but given that climate change is leading to both increased daily temperatures and increased severity of winter weather events, understanding thermal plasticity is also key to predicting how ectotherm ecology and evolution might change (5–7).

A substantial and growing body of literature details how insect RCH and cold acclimation are mediated by a variety of physiological, genetic, and biochemical changes, many of which are tied to ionoregulatory balance (8–17). There is also evidence that absolute temperature information encoded by cold sensors drives changes in sleep and wakefulness, and that shutdown in the central nervous system (via spreading depolarization) is coincident with cold acclimation (18, 19). Yet, it remains largely unknown what proximally activates cold acclimation, or to what degree peripheral sensory neurons might function in this capacity – long-standing questions in ectotherm biology (3, 20, 21). This is especially important for larvae, which are typically more sensitive to cold shocks than adults (2).

Given the suite of genetic tools available in *Drosophila*, and an emerging understanding of *Drosophila* larval cold nociception, the fruit fly larva constitutes a useful organism for investigating these unknowns. In response to noxious cold (≤10°C) *D. melanogaster* larvae primarily execute a highly stereotyped, bilateral contraction (CT) response, where the head and tail tuck toward the midline (22). This behavior is triggered by the activation of cold nociceptors innervating the barrier epidermis. In larvae, the primary cold nociceptors have been demonstrated to be Class III (CIII) dendritic arborization (da) peripheral sensory neurons, with Class II da (CII) and chordotonal (Ch) neurons also functioning in the noxious cold-sensing neural ensemble (22, 23). In contrast, innocuous cool sensing takes place among Ch neurons and thermosensors in the dorsal and terminal organ ganglia; these sensors primarily inform thermotaxis, which is undoubtedly an important part of maintaining optimal temperature (24–26).

Given that CT results in a reduced surface area to volume ratio—which is a common strategy for keeping warm (27–31)—cold nociception appears, *prima facia*, to be protective. However, it is unknown how widespread cold nociception is among drosophilids, to what degree cold nociception protects against cold shocks, or how cold nociception might relate to cold acclimation.

Herein, we more completely assess cold nociceptive behavior among drosophilid larvae, describing cold-evoked behaviors in 11 species with known differences in their cold tolerance (32–34). We additionally test the hypotheses that cold nociceptors are necessary and sufficient to cold acclimation. Making use of cold-behavior and cold-shock assays, optogenetics, electrophysiology, and methods to genetically disrupt neural transmission, we demonstrate that: (1) Although behavioral programs differ between drosophilid species, cold nociception is broadly present and CT is the predominant cold-evoked behavior; (2) species within the repleta group perform a unique, highly stereotyped, cold-evoked behavior we termed the spiracle extension response (SER); (3) in *D. melanogaster*, silencing CIII nociceptors results in an inability to cold acclimate; (4) cold acclimation is coincident with nociceptor sensitization; and (5) optogenetic activation of CIII nociceptors, *sans* cold, is sufficient for driving cold acclimation.

## Results

### Cold nociception is widespread among drosophilid larvae

We have previously demonstrated that *D. melanogaster* larvae primarily respond to noxious cold by performing a bilateral contraction (CT) along the head-to-tail axis (22). In order to elucidate a broader understanding of drosophilid cold nociception, we assessed cold-evoked behaviors (0-16°C) among 11 drosophilid species with known differences in distribution and cold tolerance (**Figure 1A-B**) (32–34).

**Figure 1.**
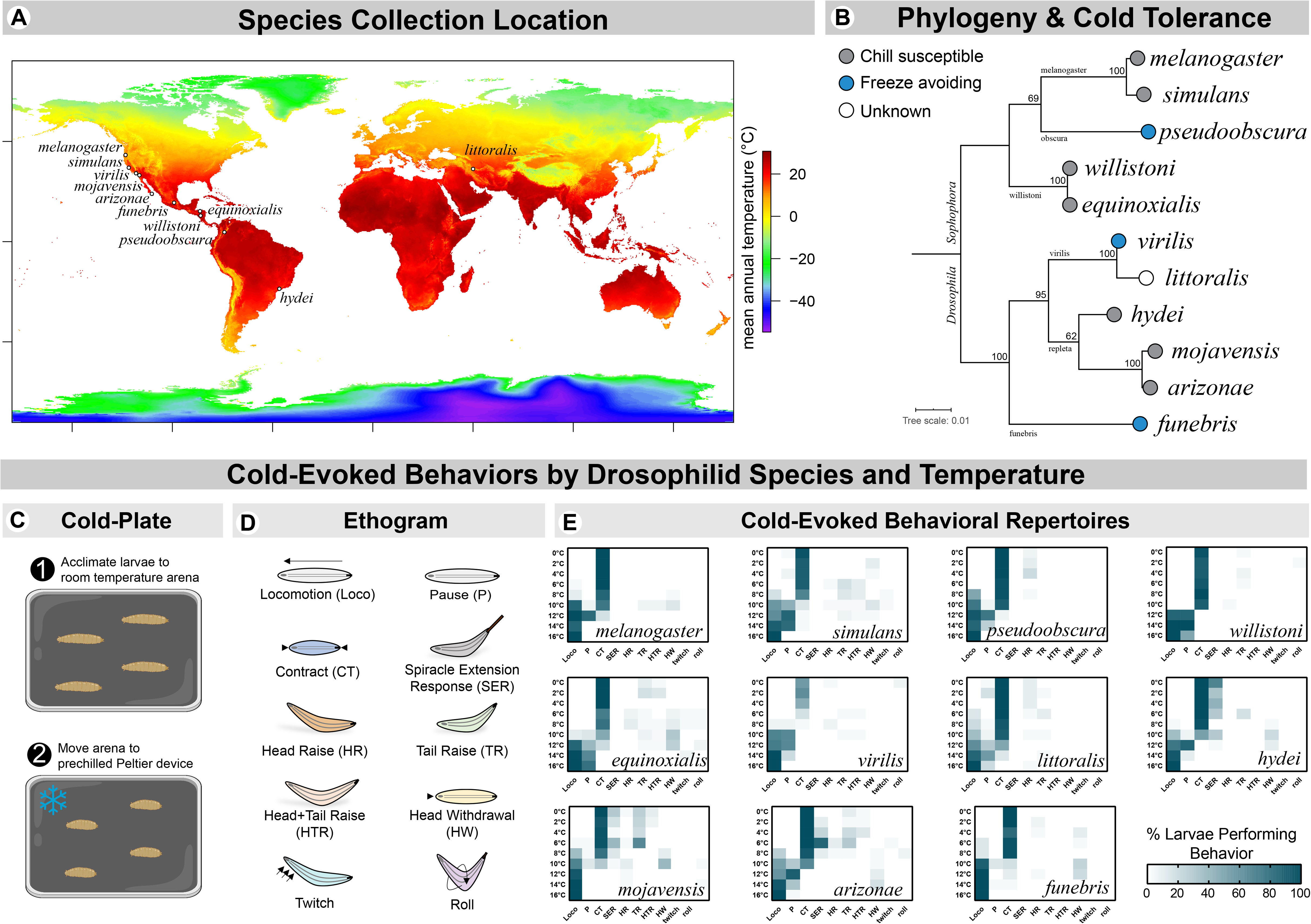
*Drosophila* species native to a variety of thermal environments and with known differences in cold tolerance were chosen for behavioral analysis. (**A**) Map of species stock collection location (provided by the *Drosophila* Species Stock Center) mapped on mean annual temperature, as extracted from climate data retrieved from WorldClim (https://www.worldclim.org) (56). (**B**) Maximum likelihood phylogeny with ultrafast bootstrapping (at nodes). This tree was generated using concatenated and aligned *CoI*, *CoII*, and *Adh* sequences (**Table S1**). Most species chosen are chill susceptible (die above freezing point) while 3 are freeze avoiding (able to maintain liquid body fluids below 0°C). Cold tolerances from (34). (**C**) Outline of the previously developed cold-plate behavioral assay (22, 57). Subject behaviors were recorded for 30s after application of cold. (**D-E**) Ethogram and heatmap representations of full cold-evoked behavioral repertoires by species and temperature. The predominant cold-evoked response among drosophilid larvae is a bilateral, head-and-tail contraction (CT) behavior. Among repleta group species, an additional, highly stereotyped behavior termed the spiracle extension response (SER) is present at low temperatures. CT was typically transient; locomotion resumed at higher temperatures (at and above appx. 8°C, depending on species), and lasting, relaxed paralysis occurred at lower temperatures (appx. 0-10°C, depending on species). Larvae performed a variety of other behaviors at relatively low frequencies, with no obvious patterns. N=2,970; n=30 for each condition.

Using the previously developed cold plate assay (22) (**Figure 1C**) we observed 9 distinct, cold-evoked behaviors, with each species exhibiting a slightly different behavioral repertoire (**Figure 1D-E** and **Tables S2-S12**). All species tested performed the CT response (**Figure 1E and Figure 2A**), yet larvae typically failed to remain contracted for the 30 second behavioral analysis. The majority of larvae either ceased CT and resumed locomotion (at higher temperatures; at and above ~8°C, depending on species) or ceased CT and were paralyzed by the cold (at lower temperatures; ~0-10°C, depending on species). It therefore appears unlikely that the CT behavior itself would protect larvae from long-term noxious cold exposure, at least not by solely reducing the animal’s surface-area-to-volume ratio.

**Figure 2.**
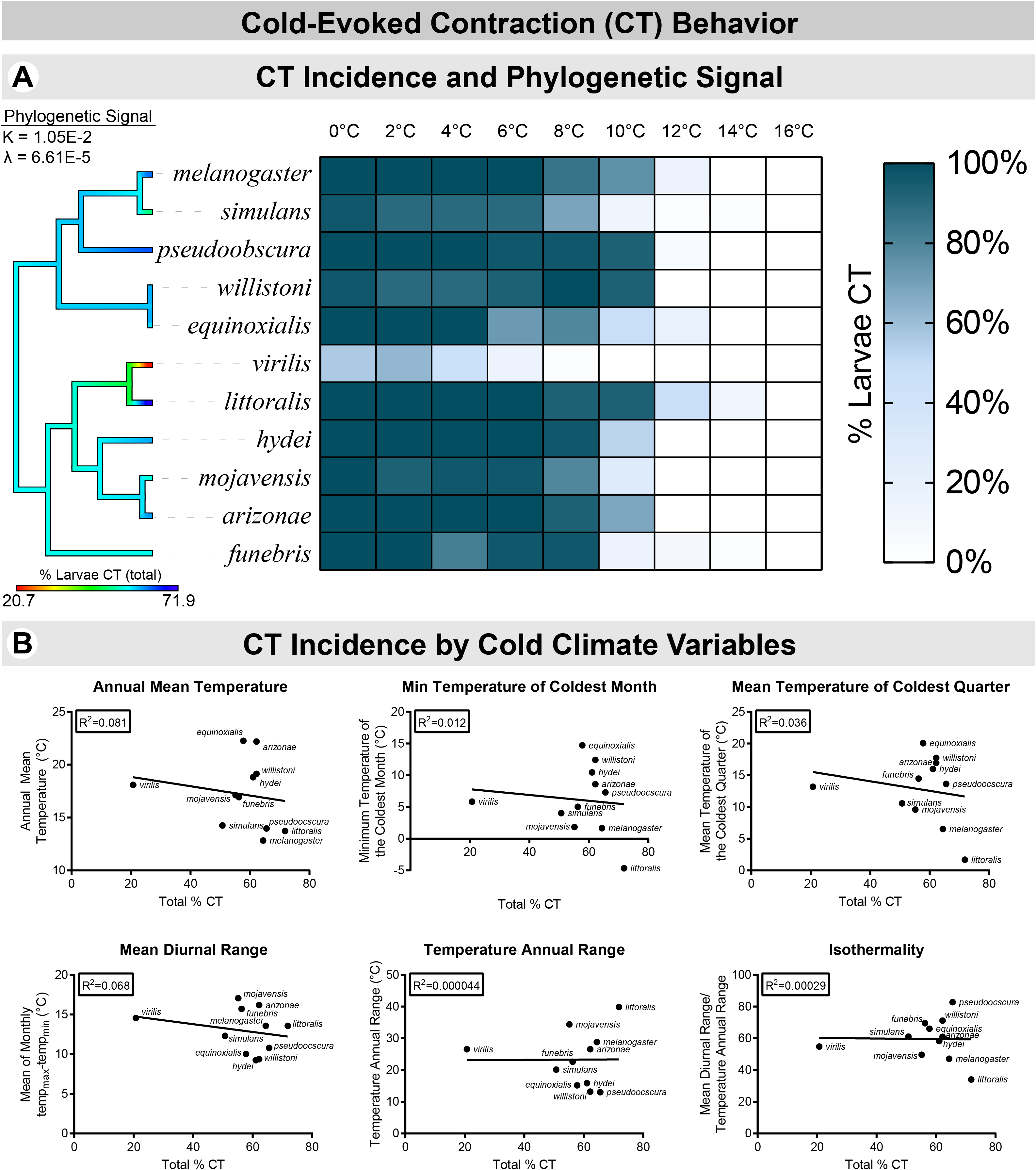
(**A**) Heatmap representation of % CT by species and temperature. All species CT, but with slightly different cold sensitivities. N=2,970; n=30 for each condition. % of larvae which performed CT out of the total intraspecies population was color-mapped to the tree by maximum likelihood ancestral state estimation (color at tip indicates value for that species). Blomberg’s K (35) and Pagel’s λ (36) show no evidence of phylogenetic signal. (**B**) Correlation analyses of % larvae CT (total) by climate variables retrieved from WorldClim. No significant correlations were found.

We predicted that, if CT was adaptive, there might be evidence of trait lability and/or of thermal environment placing selective pressure on its evolution. We assessed this by measuring phylogenetic signal and investigating possible environmental correlates of CT behavior. Computed measures of phylogenetic signal—Blomberg’s K (35) and Pagel’s λ (36)—show no evidence of phylogenetic signal for the incidence of CT (% CT in the entire intraspecies population tested; **Figure 2A**). Neither measure was statistically different from 0 (K=1.05E-1 and λ=6.61E-5), indicating that closely related species were no more similar in their CT incidences than they were to randomly selected, more distantly related species. This has typically been interpreted as trait lability, an indication of rapid evolution, or that the measured trait is not evolving as a result of constant-rate genetic drift (37–39). The most conservative interpretation, however, is simply that interspecies variation in CT incidence is not consistent with a model of evolution built on Brownian motion. However, we found no correlations between CT incidence and a variety of cold-related climate variables (**Figure 2B**), possibly indicating that the behavior is not evolving directly in response to differences in cold climate.

In addition to CT, a second notable behavior we termed the spiracle extension response (SER) was observed (**Figure 1D-E** and **Figure 3**). In larvae, the posterior spiracles are part of a snorkel-like respiratory organ with the ability to extend and contract, allowing larvae to respirate while mostly submerged in semi-liquid media (40). In response to noxious cold, larvae from the repleta group performed a highly stereotyped behavior wherein the posterior stigmatophore—which contains the spiracular chamber—rapidly and greatly extended from the contracted state (**Figure 3A**). This behavior typically occurred post-CT and was accompanied by a robust tail-raise behavior. This behavior does not seem to be solely initiated by direct, chill-induced contractions (41) in the musculature controlling the stigmatophore, as segmental cold stimulation across the body also resulted in the SER (**Figure 3B**). While CT likely emerged in a common drosophilid ancestor, SER likely emerged within the repleta group (supported by ancestral state reconstruction; **Figure 3C**).

**Figure 3.**
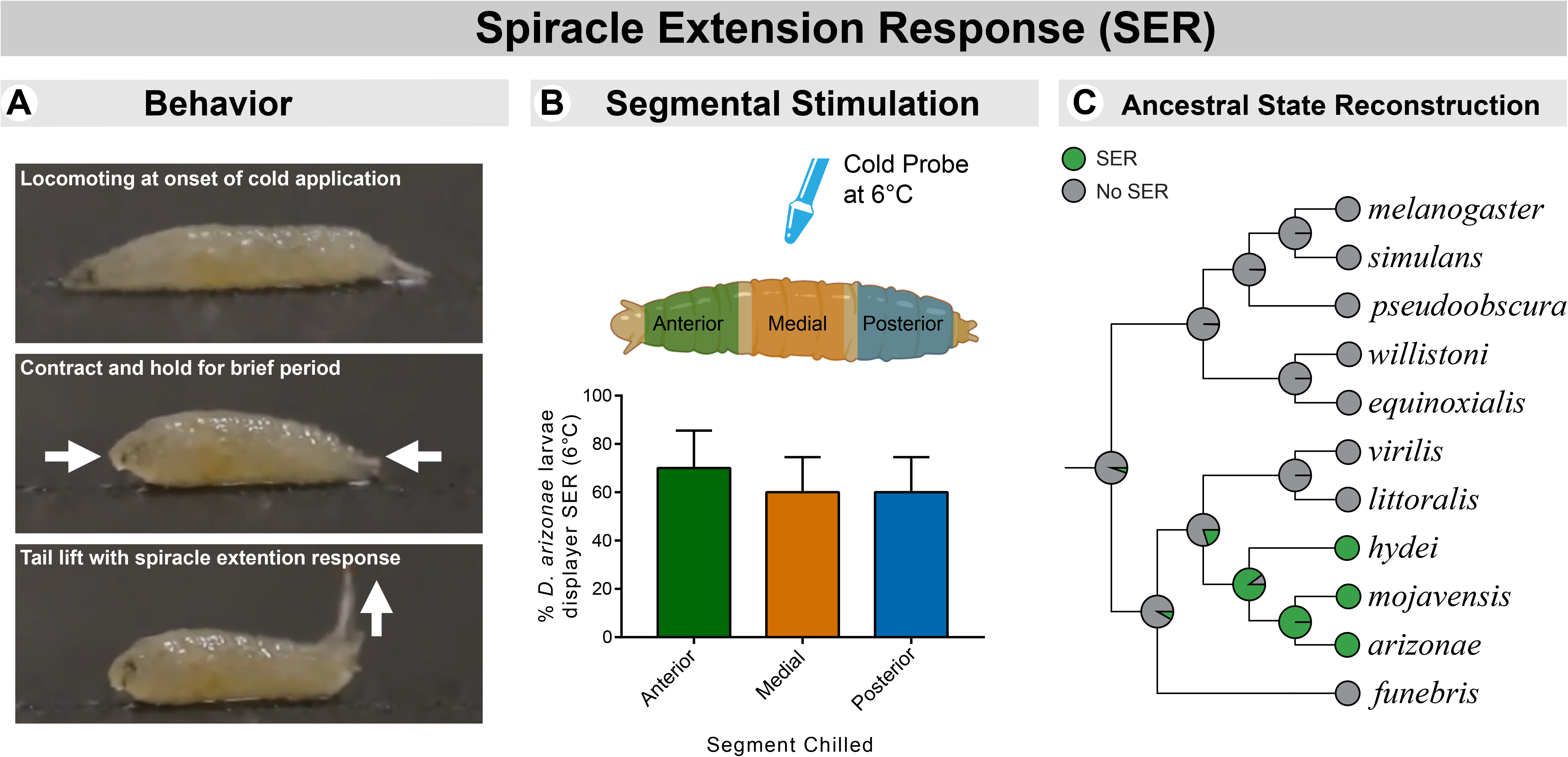
The SER is a highly stereotyped behavior that likely evolved specifically within the repleta group. (**A**) Description of cold-evoked SER, with images (subject, *D. arizonae*). (**B**) % SER following segmental cold stimulation via cold-probe assay. N=30, n=10. (**C**) Ancestral state reconstruction by single-rate continuous-time Markov chain model indicates that the SER behavior most likely evolved in a common repleta ancestor, after the group split from other drosophilids.

### Silencing CIII cold nociceptors inhibits cold acclimation

Given the widespread presence of cold nociception among drosophilids, we next questioned whether cold nociception plays a role in protecting larvae from noxious cold.

We first tested the hypothesis that cold nociception is a necessary component of cold acclimation. The *GAL4-UAS* system was used to drive the expression of the light chain of tetanus toxin (TNT) in targeted cold-sensing neural populations (CII, CIII, Ch, CII+CIII, and CIII+Ch), thereby selectively silencing neural transmission. TNT expression in cold nociceptors has been previously shown to severely inhibit the cold-evoked CT behavioral response (22).

Transgenic *D. melanogaster* were raised at 24°C and then incubated for 48 hours at either 24°C or 10°C, the latter to drive cold acclimation. After incubation, 3^rd^ instar larvae were cold-shocked at 0°C for 60 minutes by a modified cold-plate assay. Survival was assessed by the proportion of animals that eclosed as adults (**Figure 4A**). In order to control for any baseline differences in cold tolerance between transgenic strains, cold tolerance was also assessed in subjects expressing an inactive form of TNT (impaired TNT; *TNT ^IMP^*) in the same *GAL4* genetic backgrounds.

**Figure 4.**
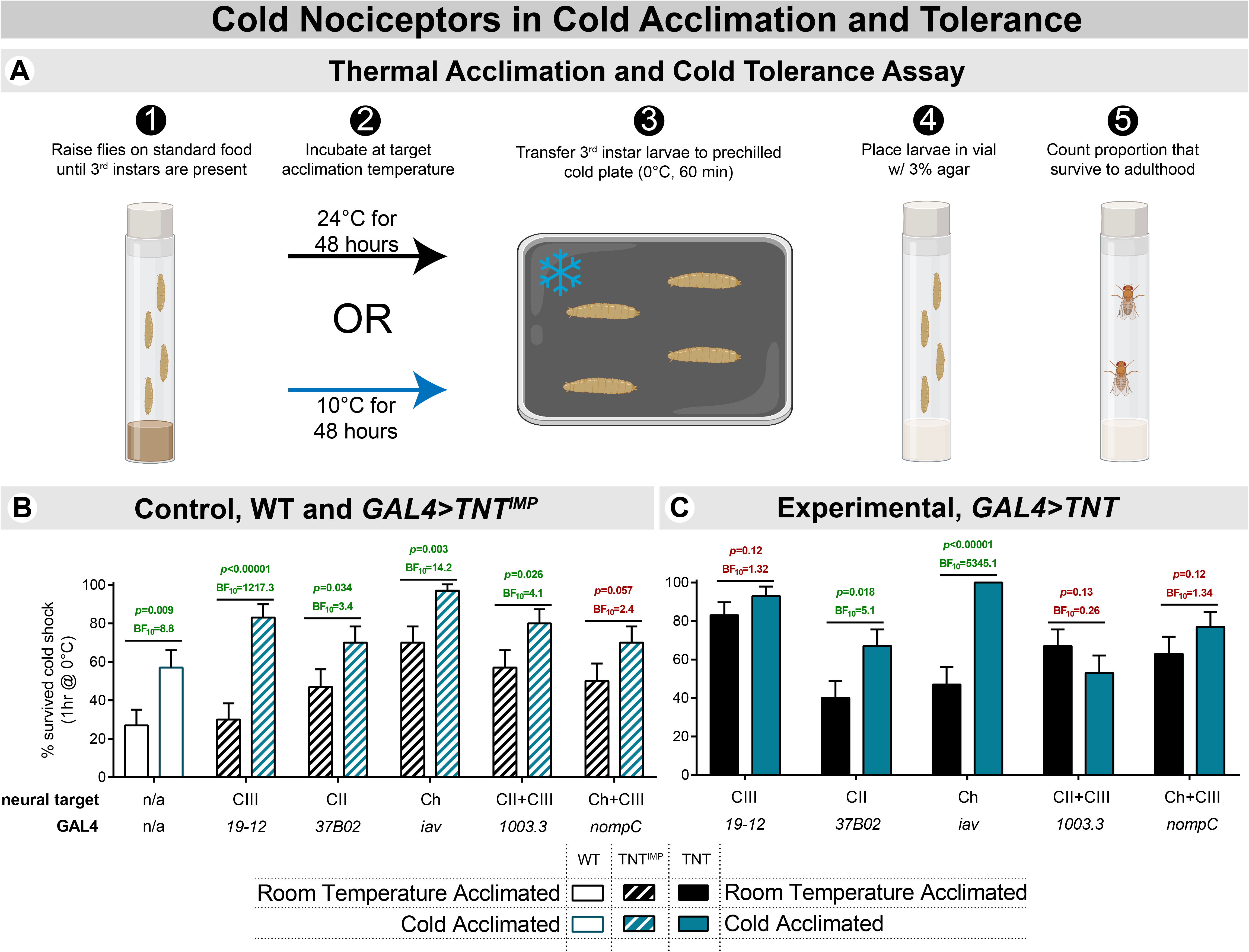
Class III nociceptors are necessary for cold acclimation. (**A**) Outline of the approach used for thermal acclimation and cold tolerance assessment. *D. melanogaster* larvae were acclimated to either 24 or 10°C and subjected to 0°C cold shock for 60 minutes. The percentage of animals which successfully eclosed constituted the % survival rate. (**B**) Control (*w ^1118^* and *GAL4>TNT ^IMP^*) larvae showed some baseline differences in cold tolerance, but showed increased cold tolerance following cold acclimation, regardless. N=360; n=30 in each condition. (**C**) Silencing CIII neurons via active tetanus toxin (TNT) resulted in an inability to cold acclimate; cold acclimated larvae had similar cold tolerance to room temperature acclimated larvae. Silencing CII and Chordotonal (Ch) neurons independently had no effect on cold acclimating capacity. Silencing CIII neurons independently resulted in an increase in baseline cold tolerance, but this was not apparently present when combinatorially silencing CIII with CII/Ch. N=300; n=30 in each condition.

In accordance with previous studies (42–47), cold acclimated control animals (*w ^1118^* and *TNT ^IMP^*) were more cold tolerant than counterparts raised at room temperature (**Figure 5B**). Although the *nompC* control (CIII+Ch) condition was not significantly or substantially different, Bayesian statistics indicate that the hypothesis that they cold acclimate is ~2.4 times more likely than the null hypothesis.

**Figure 5.**
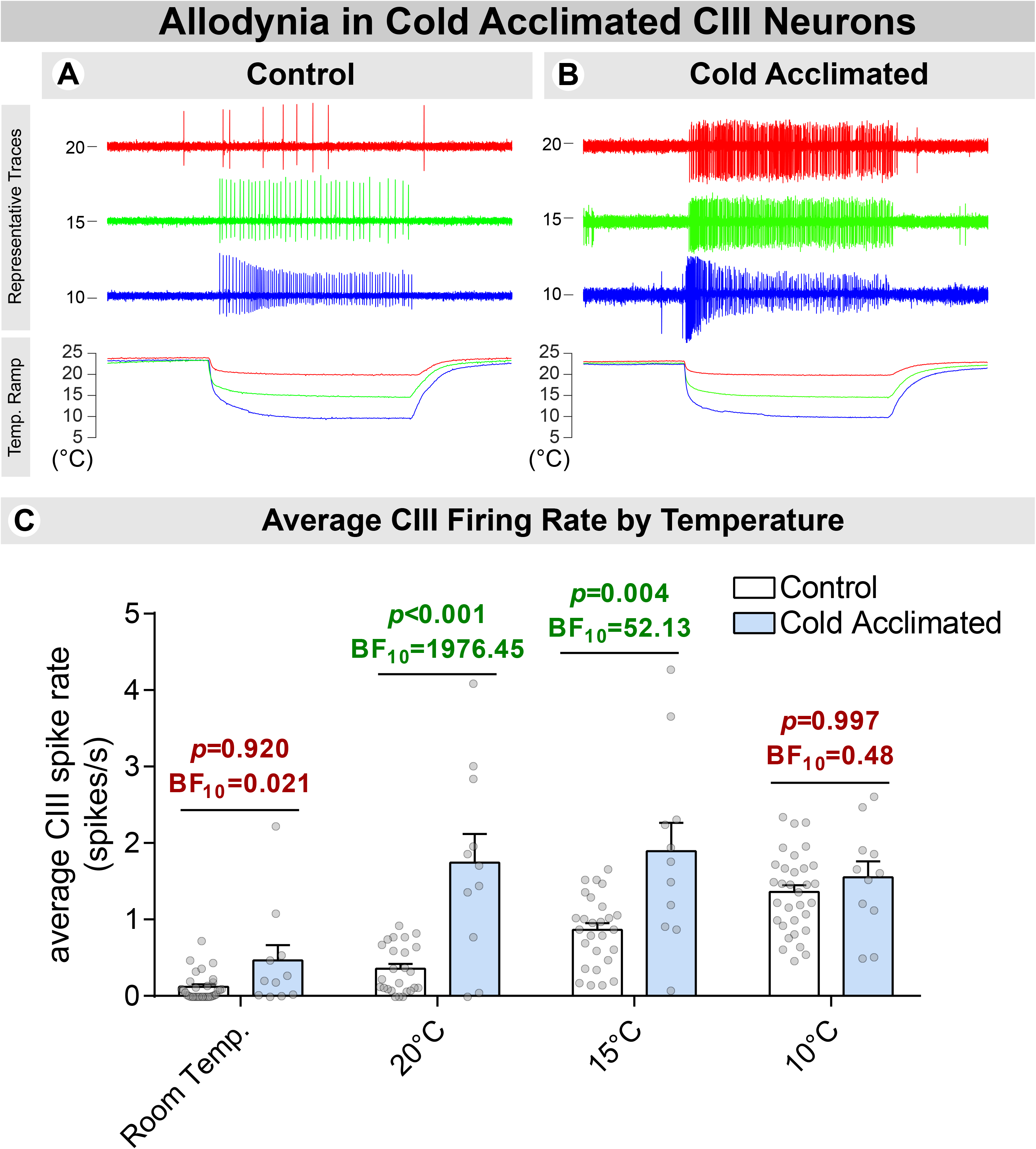
Cold acclimation results in nociceptive sensitization. (**A-B**) Representative traces of extracellular CIII recordings in control and cold acclimated larvae. Neural activity was assessed at room temperature, and during/following ramps to 20, 15, and 10°C. N=43; control, n=32; cold acclimated, n=11. (**C**) Average spike frequency of CIII neurons by temperature. CIII neurons in cold acclimated larvae were more sensitive to temperature drops to 20 and 15°C.

In contrast, silencing CIII neurons resulted in an inability to cold acclimate, indicating that CIII cold nociceptive neurons are necessary for cold acclimation **(Figure 5C**). Silencing CII and Ch neurons alone had no obvious effects – larvae were still capable of cold acclimating. Surprisingly, silencing CIII neurons alone (*19-12 GAL4*) resulted in increased cold tolerance in room temperature-acclimated larvae, mimicking survival rates of cold-acclimated larvae.

Combinatorially silencing CIII with CII or Ch neurons resulted in an inability to cold acclimate, but did not obviously increase baseline cold tolerance.

Given that silencing CIII neurons via 3 different *GAL4*s (*19-12*, *1003.3*, and *nompC*) resulted in an inability to cold acclimate, the preponderance of evidence suggests that CIII neurons are necessary for cold acclimation. Moreover, that single-class and CIII-combinatorial silencing of CII and Ch (via *1003.3* and *nompC GAL4*s) doesn’t result in an obvious phenotype in room temperature acclimated animals, these results suggest that CII and Ch neurons are not necessary for cold acclimating capacity, but that they may play some role in modulating cold tolerance. This is consistent with CII and Ch neurons functioning in cold nociception, but not in an independently necessary capacity (22, 23).

We next asked how cold acclimation impacts CIII function by recording cold-evoked CIII activity following cold acclimation. Larvae were raised at 24°C and then incubated for 48 hours at either 24°C or 10°C, as above. Cold-evoked electrical activity was recorded from CIII neurons in live, filleted larvae; CIII neurons were identified by *GAL4-UAS*-mediated GFP labeling (*GAL4^19-12^>UAS-mCD8::GFP*).

Electrophysiological recordings of CIII neurons reveal that cold acclimation results in CIII hypersensitization; on average, CIII neurons in cold acclimated larvae responded more strongly to temperature drops to 20°C and 15°C (**Figure 5**). Firing rates were increased and sustained at 20°C and 15°C, and although mean firing rate was not significantly or substantially increased at 10°C (**Figure 5C**), we observed an increased firing rate at the onset of chilling (first 10 seconds), with activity quickly returning to control levels (**Figure S1**).

### Developmental optogenetic-mediated CIII activation increases cold tolerance

As CIII nociceptors were necessary for cold acclimation, we next tested the hypothesis that CIII nociceptors drive cold acclimation by optogenetically activating CIII neurons *sans* cold and assessing cold tolerance.

The *GAL4-UAS* system was used to drive expression of an engineered channelrhodopsin (ChETA)—a light-gated cation channel—in CIII neurons. Transgenic *D. melanogaster* were raised on food containing all-*trans*-retinol (ATR+, which is a necessary for the activity of ChETA) or standard food (ATR-, control condition). Larvae were housed in Petri dishes with 2.5mL of food spread extremely thin over the base. 48 hours prior to assessing cold tolerance, dishes were transferred to a custom built OptoBox which houses 12 independently operated chambers capable of delivering blue light at preset intervals. Blue light (~470nm) was delivered for 5s every 5 minutes, a paradigm previously used for developmental activation of CIV nociceptors (48). After 48 hours, larvae were cold-shocked at 0°C for 60 minutes (**Figure 6A**).

**Figure 6.**
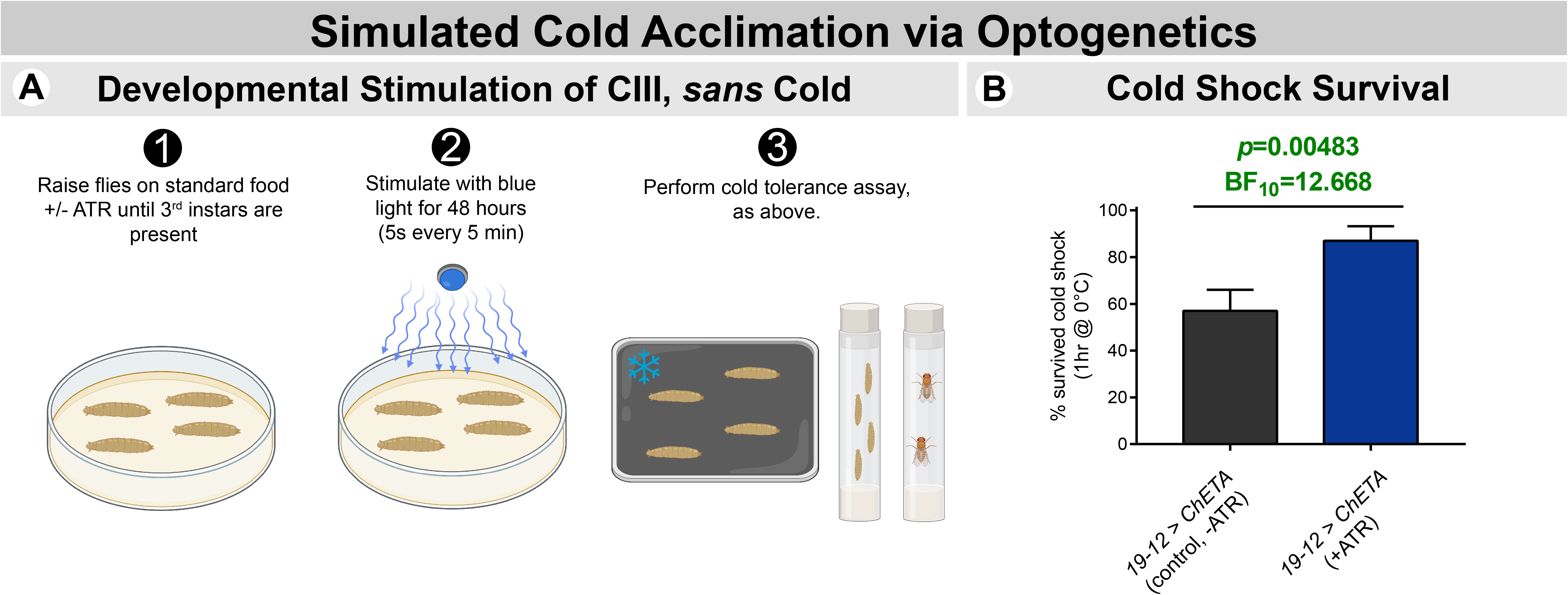
(**A**) Outline of optogenetic acclimation and cold tolerance assay. Larvae expressing ChETA in CIII neurons were raised on food + or – ATR and then subjected to 48 hours of blue light exposure (5s every 5 min). (**B**) Activating CIII neurons optogenetically, *sans cold*, resulted in larvae with higher cold tolerance. N=60; n=30 in each condition.

Long-term, repeated activation of CIII neurons improved survival rates following cold shock, indicating that the activation of cold nociceptors is sufficient for driving cold acclimation (**Figure 6B**).

## Discussion

Herein, we have found some conflicting evidence concerning the adaptive nature of the CT response. The CT response is widespread, occurs primarily at low, potentially harmful temperatures, results in a reduced surface-area-to-volume ratio, and is conserved from a common ancestor, collectively indicating that the behavior might be adaptive. The lack of phylogenetic signal we detected may also be consistent with this, as values close to 0 may indicate trait lability, perhaps suggesting that cold-evoked behavior has rapidly adapted as drosophilids have entered new thermal environments.

Several points are worth considering, however. Firstly, we found no evidence that CT incidence was correlated with any particular cold-climate variable. Secondly, we used a relatively small sample of species, and only a single strain per species, which may lead to a degree of uncertainty in both our assessments of intraspecies behavior and phylogenetic signal. It remains commonplace and acceptable to use single strains in comparative *Drosophila* studies (32, 34, 49, 50), however some studies have shown substantial differences in a variety of metrics within species, across strains (51–54). Our work here also evidences this, given that different *GAL4* genetic backgrounds appeared to have different baseline cold tolerances (**Figure 4B**). Thirdly, low phylogenetic signal, despite being an indication of trait lability, does not give evidence of any specific type of selection. The absence of phylogenetic signal can therefore be interpreted several different ways; for example, it may indicate rapid adaptation into specific niches, stabilizing selection, and/or evolution under constant functional constraint (37–39). Fourthly, we observed that cold-evoked behaviors are highly transient – at low temperatures larvae eventually enter a state of flaccid paralysis. Finally, we observed no obvious baseline collapse in cold-tolerance as a result of impairing cold nociception via TNT – in fact, one condition resulted in dramatically increased cold tolerance (**Figure 4C**; *19-12 GAL4*).

With all of these considerations in mind, we do not believe there is a strong argument to be made that CT, or any noxious cold-evoked behavior, has long-term protective qualities in and of itself. Although further work is required to better clarify the evolutionary history and plasticity of insect cold nociceptive behavior, what remains clear is that drosophilid CT behavior is widespread and inherited from a common ancestor. Despite it possibly not being protective on its own, CT is an easily evoked, highly identifiable behavior under the control of multimodal nociceptors (22, 23). It therefore remains a useful tool for studying potentially generalizable molecular, cellular, and systems-level mechanisms of nociception and sensory multimodality.

In contrast, cold nociception is necessary for cold acclimation; silencing CIII nociceptors results in an inability to cold acclimate. Moreover, CIII activation appears sufficient for inducing cold acclimation, as activating CIII neurons optogenetically leads to increased cold tolerance. Whether or not the nervous system plays a role in thermal acclimation has been a long-standing question in ectotherm biology (3, 20, 21). These findings provide evidence that the sensory nervous system is critical to cold acclimation. Given the high genetic tractability of *Drosophila*, this opens several avenues for future mechanistic studies.

Interestingly, cold acclimation also resulted in nociceptor sensitization, arguably a form of allodynia (the state of experiencing pain in response to less-than-noxious stimuli); we observed increased cold sensitivity in the CIII neurons of cold-acclimated larvae. One possible hypothesis is that cold acclimation induces a state of hyper-vigilance in response to chilling, therefore better preparing larvae for cold shocks. Given that cold acclimation is likely the result of long-term changes in physiology and gene expression (13, 16), it’s unclear how this hyper-vigilance might lead to increased cold tolerance, in and of itself. However, the presence of cold-induced hypersensitivity and the coincident increase in cold tolerance may be consistent with the hypothesis that nociceptive hypersensitization provides survival advantage, a hypothesis formulated by Crook, Walter, and colleagues, who observed that injury-induced nociceptor sensitization results in decreased predation risk in squid (55). Whether injury plays any role in cold acclimation is a subject which requires further study. Additional studies are also required in order to elucidate whether or not cold tolerance is necessarily tied to sensitization, or if nociceptive sensitization is simply a coincident phenomenon.

In summary, we have shown that cold nociception is widespread among drosophilids and that *D. melanogaster* CIII neurons are necessary and sufficient components of cold acclimation. The results of these experiments suggest that insect cold acclimation has a peripheral neural basis in cold nociception. These findings will inform future studies which will further our understanding of thermal plasticity, particularly in the context of rapidly changing climate dynamics.

## Materials & Methods

### Animals

All *Drosophila* stocks were maintained at 24°C. The *OregonR* (*ORR*) and *w^1118^* strains were used to assess wild-type (WT) behavior and cold tolerance, respectively, in *D. melanogaster*. Transgenic *D. melanogaster* strains included: *GAL4^GMR37B02^* (CII neurons); *GAL4^19-12^* (CIII neurons); *GAL4^iav^* (Ch neurons); *GAL4^1003.3^* (CII and CIII neurons); *GAL4^nompC^* (CIII and Ch neurons, BDSC #36361); *UAS-TeTxLC* (active tetanus toxin light chain, TNT, BDSC #28837); *UAS-TNT^IMP^* (impaired tetanus toxin light chain, BDSC #28840); *UAS-mCD8::GFP* (CD8-mediated membrane targeting of GFP, BDSC #5130); and *UAS-ChETA::YFP* (engineered derivative of Channelrhodopsin2, BDSC #36495).

Other drosophilid species were obtained from The National *Drosophila* Species Stock Center at Cornell University: *Drosophila arizonae* (#15081-1271.39); *Drosophila equinoxialis* (#14030-0741.00); *Drosophila funebris* (#15120-1911.01); *Drosophila hydei* (#15085-1641.04); *Drosophila willistoni* (#14030-0811.00); *Drosophila virilis* (#15010-1051.00); *Drosophila simulans* (#14021-0251.292); *Drosophila pseudoobscura* (#14011-0121.35); *Drosophila mojavensis* (#15081-1352.47). Species distribution on map of annual mean temperature was generated in R using climate data retrieved from WorldClim (https://www.worldclim.org) (56) and location data provided by the *Drosophila* Species Stock Center.

### Behavior Assays

Cold-evoked behaviors were assessed using the previously developed cold-plate and cold-probe assays (22, 57, 58). For cold-plate, wandering 3^rd^ instar larvae were acclimated to a room-temperature aluminum arena. The arena was then transferred to a prechilled Peltier device (TE Technologies, CP0031) under the control of a thermoelectric temperature controller (TE Technologies, TC-48-20). Behavior was recorded from above for 30 seconds and assessed qualitatively *post hoc*. For cold-probe, the cold probe apparatus (ProDev Engineering) was chilled to 6°C, and then placed at a 45° angle upon a posterior, medial, or anterior segment of the larva, and held for 5 seconds; behavior was scored qualitatively at the time of the assay.

### Cold Survival Assay

Larval cold tolerance was assessed using a modified cold-plate assay. *Drosophila* populations were raised at 24°C until 3^rd^ instar larvae were present. The entire population was then incubated for 48 hours at 24°C or 10°C. After the incubation, wandering 3^rd^ instar larvae were collected using a small paint brush, transferred to a room-temperature aluminum arena, and allowed to acclimate until locomotion resumed. To deliver cold shock, the arena was transferred to a Peltier device prechilled to 0°C. Larvae were cold-shocked for 60 minutes, then removed from the arena and placed in a new *Drosophila* vial. As wandering 3^rd^ instar larvae are post-feeding, cold-shocked larvae were housed in vials containing 10mL of 3% agar instead of food (to prevent desiccation and confounds due to food molding). Larvae were cold-shocked immediately after incubation and the remaining populations were discarded. Survival was assessed by counting the eventual number of adults eclosed as compared to the number of larvae originally placed in the vial.

### Optogenetics

*GAL4^19-12^>UAS-ChETA::YFP* larvae were raised in Petri dishes, in constant darkness, at 24°C. As channelrhodopsins require all-*trans*-retinol (ATR) to function, larvae with identical genotypes were either raised on food containing ATR (ATR+, 1.5mM) or standard food (ATR-, as control). When 3^rd^ instar larvae were present in the dish, the dish was transferred to a custom-built OptoBox (48), where high intensity blue light was automatically delivered at a frequency of 5 seconds every 5 minutes, for 48 hours. The OptoBox consists of a plastic storage bin segmented into 12 independent test chambers, each equipped with a blue LED with an 80° lens (RapidLED, CREe XP-E2) under the control of an Arduino microcontroller (Arduino, Uno Rev3). After 48 hours, the dishes were then removed from the OptoBox and wandering 3^rd^ instar larvae were used in the cold survival assay detailed above. Larvae were shocked immediately following removal from the OptoBox, and the remaining populations were discarded.

### Phylogenetics

Amino acid sequences (**Table S1**) for Cytochome Oxidase I/II (mt:*CoI*/*CoII*) and alcohol dehydrogenase (*Adh*) were aligned via MAFFT, using default settings. As the repleta group has two copies of *Adh*, we used *Adh1* for these species. Phylogenetic trees were generated via IQ-Tree by the maximum likelihood approach, using an LG+F+R2 substitution model (as determined by ModelFinder). Branch support was calculated by ultrafast bootstrapping (UFboot, 2000 replicates). Trees were visualized using iTOL, R, and Adobe Illustrator.

All additional phylogenetic analyses were performed in R using the phytools package. SER ancestral state reconstruction was performed using a single-rate continuous-time Markov chain model; pie-charts at interior nodes of the presented cladogram represent empirical Bayesian posterior probabilities. % larvae CT (out of total population) ancestral states were mapped to an ultrametric phylogeny by maximum likelihood via the chronos and contMap functions.

### Electrophysiology

Demuscled fillet preparations were made from *GAL4^19-12^>UAS-mCD8::GFP* larvae housed in Petri dishes lined with Sylgard 184 (Dow Corning), which were filled and constantly superfused with HL-3 saline. Extracellular recordings were made with a pipette (tip diameter, 5–10 μm) connected to the headstage of a patch-clamp amplifier (Multiclamp 700A, Molecular Devices). Gentle suction was applied to draw the soma and a small portion of neurite into the pipette. The amplifier was set to voltage-clamp mode to record neuronal spikes. The output signals from the amplifier were digitized at a sampling frequency of 10 kHz using a Micro1401 A/D converter (Cambridge Electronic Design) and acquired into a laptop computer running Windows 10 with Spike2 software v. 8 (Cambridge Electronic Design). To apply a low temperature stimulation, saline was passed through an SC-20 in-line solution cooler (Warner Instruments) connected to a CL-100 temperature controller (Warner Instruments). Average spike frequency was measured during baseline room-temperature conditions (60 seconds) and during superfusion of chilled saline (60 seconds at 20, 15, or 10°C).

### Statistics

Due to a growing call for statistical analyses that do not rely on *p*-values (59–67), differences in mean and population proportions were analyzed using both traditional frequentist statistics and Bayesian alternatives. Nonlinear regression analyses were performed in GraphPad PRISM, z-tests were performed in Microsoft Excel, and all other analyses were performed in JASP using default prior probability distributions (68). Population proportions are presented as % ± standard error of the proportion (SEP); differences in proportion were assessed by one-tailed z-test with the Benjamini-Hochberg procedure (false discovery rate at 0.05) and the Bayesian A/B Test. All other measures are presented as mean ± standard error of the mean (SEM); differences were assessed by frequentist ANOVA with a Scheffé correction for multiple comparisons, and the Bayesian ANOVA with multiple comparisons corrected by the Westfall-Johnson-Utts method (69).

For Bayesian analyses, degree of support for rejecting the null hypothesis was inferred by computed Bayes Factors (BF), via the method originally proposed by Jeffreys (1961): “no evidence” (BF_10_ < 1); “barely worth mentioning” (BF_10_: 1-3); “substantial” (BF_10_: 3-10); “strong” (BF_10_: 10-30); “very strong” (BF_10_: 30-100); and “decisive (BF_10_ > 100).

Histograms and heatmaps were generated using GraphPad PRISM (GraphPad Software, La Jolla, California, USA). In statistics bars, green text indicates statistical significance (α=0.05, frequentist) or substantial evidence in favor of the alternative hypothesis (BF_10_ > 3, Bayesian). Red text indicates no statistical significance (frequentist) or less than substantial evidence in favor of the alternative hypothesis (Bayesian). Population sizes (N) are indicated in figure legends. n=30 for each condition unless otherwise stated.

## Funding

This work is supported by NIH R01 NS115209-01 (to DNC). NJH was supported by NIH F31 NS117087-01, a GSU Brains & Behavior Fellowship, and a Kenneth W. and Georganne F. Honeycutt Fellowship. JML was supported by a GSU Brains & Behavior Fellowship.

## Author Contributions

Conceptualization, NJH; Methodology, NJH, JML, AS, KJD, and DNC; Cold-plate assays, NJH, JML, TRG, and MNB; Survival assays, NJH; OptoBox design and construction, NJH and KJD; Optogenetics, NJH; Electrophysiology, AS; Photography, TRG; Phylogenetics and associated analyses, NJH; Statistics and other formal analyses, NJH; Writing – Original Draft, NJH; Writing – Review & Editing, NJH, JML, AS, TRG, MNB, KJD, and DNC; Visualization, NJH and AS; Supervision, DNC; Funding Acquisition, NJH and DNC.

## Acknowledgments

We thank Dr. Bing Ye (University of Michigan) for providing materials lists, wiring diagrams, and advice which contributed to the design and construction of the OptoBox, and Dr. Gennady Cymbalyuk (Georgia State University) for use of electrophysiological equipment. We also thank and credit BioRender, which was used to generate many of the experimental outlines and other graphics.

## Notes

### Competing Interest Statement

The authors have declared no competing interest.

