## Supplemental Information for "The evolution of cold nociception in drosophilid larvae and identification of a neural basis for cold acclimation"

Corresponding preprint in *bioRxiv*

**Nathaniel J. Himmel[1,2], Jamin M. Letcher[1,3], Akira Sakurai[1,4], Thomas R. Gray[1,5], Maggie N. Benson[1], Kevin J. Donaldson[1,6], and Daniel N. Cox[1,7]**

*1 – Neuroscience Institute, Georgia State University, Atlanta, GA, USA*
*2 – ORCid: 0000-0001-7876-6960*
*3 – ORCid: 0000-0003-3077-0615*
*4 – ORCid: 0000-0003-2858-1620*
*5 – ORCid: 0000-0002-6176-005X*
*6 – ORCid: 0000-0002-9144-3776*
*7 – ORCid: 0000-0001-9191-9212*

**Keywords:** insects, thermal plasticity, winter ecology, sensory neurons, *Drosophila*, *Sophophora*

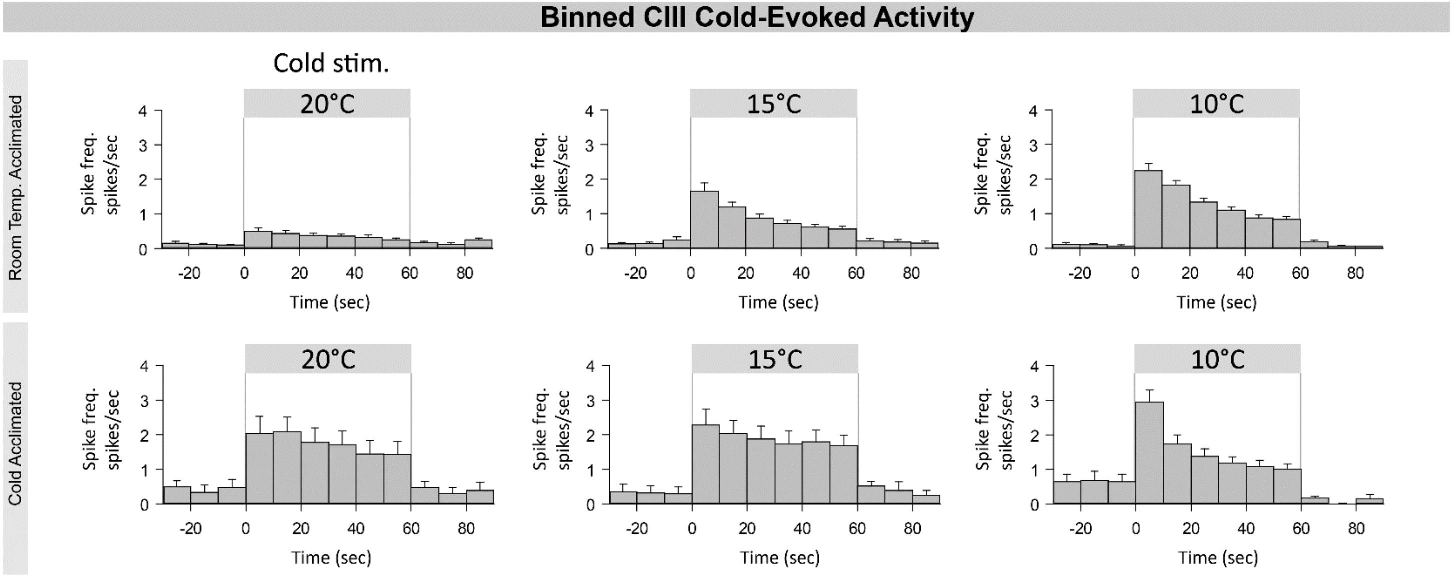


**Figure S1.** CIII neural activity by 10 second bins.  See **Figure 5**.

| | Accession Number | | |
|---|---|---|---|
| **Species** | **CoI** | **CoII** | **Adh** |
| *Drosophila melanogaster* | FBpp0100176 | FBpp0100177 | FBpp0100047 |
| *Drosophila simulans* | NP_982323 | P50253 | XP_016024999 |
| *Drosophila pseudoobscura* | YP_009742607 | KAF3759836 | XP_001357375 |
| *Drosophila willistoni* | DAA06222 | AAR05888 | XP_015033661 |
| *Drosophila equinoxialis* | AUW36081 | AAL83280 | Q9NG42 |
| *Drosophila virilis* | ADW41404 | ABD84161 | XP_002057583 |
| *Drosophila littoralis* | YP_002327403 | YP_002327404 | ABG56059 |
| *Drosophila hydei* | AFV25509 | AAD37708 | P23236 |
| *Drosophila funebris* | AHL43561 | AAM70094 | CAA73700 |
| *Drosophila mojavensis* | DAA06235 | ABD93167 | XP_002002930 |
| *Drosophila arizonae* | ABD38168 | ABD93155 | XP_017859111 |

**Table S1.** Accession numbers for sequences used to generate *Drosophila* phylogeny. Sequences from *Drosophila melanogaster* from FlyBase; all others from GenBank.

| Temp | Loco | P | CT | SER | HR | TR | HTR | HW | twitch | roll |
|---|---|---|---|---|---|---|---|---|---|---|
| | | | | | *melanogaster* | | | | | |
| 0 | 0% | 0% | 100% | 0% | 0% | 0% | 0% | 0% | 0% | 0% |
| 2 | 0% | 0% | 100% | 0% | 0% | 0% | 0% | 0% | 0% | 0% |
| 4 | 0% | 0% | 100% | 0% | 0% | 0% | 0% | 0% | 0% | 0% |
| 6 | 3% | 0% | 100% | 0% | 0% | 0% | 0% | 0% | 0% | 0% |
| 8 | 10% | 0% | 87% | 0% | 0% | 0% | 0% | 3% | 0% | 0% |
| 10 | 87% | 7% | 77% | 0% | 0% | 7% | 7% | 13% | 0% | 0% |
| 12 | 100% | 83% | 17% | 0% | 0% | 0% | 0% | 0% | 0% | 0% |
| 14 | 87% | 0% | 0% | 0% | 0% | 0% | 0% | 0% | 0% | 0% |
| 16 | 100% | 0% | 0% | 0% | 0% | 0% | 0% | 0% | 0% | 0% |

**Table S2.** *Drosophila melanogaster* cold-evoked behavioral program. Each value shows the % of larvae performing a particular behavior, at a particular temperature. See **Figure 1**.

| *simulans* | | | | | | | | | | |
|---|---|---|---|---|---|---|---|---|---|---|
| **Temp** | **Loco** | **P** | **CT** | **SER** | **HR** | **TR** | **HTR** | **HW** | **twitch** | **roll** |
| 0 | 0% | 3% | 97% | 0% | 0% | 0% | 0% | 0% | 0% | 0% |
| 2 | 0% | 0% | 90% | 0% | 0% | 0% | 0% | 10% | 0% | 0% |
| 4 | 0% | 7% | 90% | 0% | 0% | 0% | 0% | 3% | 0% | 0% |
| 6 | 0% | 0% | 90% | 0% | 3% | 17% | 10% | 0% | 0% | 0% |
| 8 | 17% | 0% | 70% | 0% | 0% | 13% | 10% | 0% | 3% | 0% |
| 10 | 59% | 45% | 14% | 0% | 3% | 3% | 3% | 14% | 3% | 0% |
| 12 | 63% | 87% | 3% | 0% | 0% | 7% | 3% | 3% | 0% | 0% |
| 14 | 90% | 83% | 3% | 0% | 0% | 0% | 0% | 0% | 0% | 0% |
| 16 | 100% | 0% | 0% | 0% | 0% | 0% | 0% | 0% | 0% | 0% |

**Table S3.** *Drosophila simulans* cold-evoked behavioral program. Each value shows the % of larvae performing a particular behavior, at a particular temperature.  See **Figure 1**.

| *virilis* | | | | | | | | | | |
|---|---|---|---|---|---|---|---|---|---|---|
| **Temp** | **Loco** | **P** | **CT** | **SER** | **HR** | **TR** | **HTR** | **HW** | **twitch** | **roll** |
| 0 | 0% | 0% | 57% | 0% | 3% | 0% | 0% | 0% | 0% | 7% |
| 2 | 0% | 0% | 63% | 0% | 0% | 0% | 0% | 0% | 0% | 0% |
| 4 | 0% | 0% | 47% | 0% | 0% | 0% | 0% | 0% | 0% | 0% |
| 6 | 0% | 0% | 17% | 0% | 0% | 3% | 3% | 0% | 0% | 0% |
| 8 | 7% | 7% | 3% | 0% | 7% | 0% | 0% | 0% | 0% | 0% |
| 10 | 70% | 67% | 0% | 0% | 0% | 3% | 3% | 0% | 0% | 0% |
| 12 | 73% | 67% | 0% | 0% | 0% | 0% | 0% | 0% | 0% | 0% |
| 14 | 100% | 20% | 0% | 0% | 0% | 0% | 0% | 0% | 0% | 0% |
| 16 | 100% | 0% | 0% | 0% | 0% | 0% | 0% | 0% | 0% | 0% |

**Table S4.** *Drosophila virilis* cold-evoked behavioral program. Each value shows the % of larvae performing a particular behavior, at a particular temperature. See **Figure 1**.

| *pseudoobscura* | | | | | | | | | | |
|---|---|---|---|---|---|---|---|---|---|---|
| **Temp** | **Loco** | **P** | **CT** | **SER** | **HR** | **TR** | **HTR** | **HW** | **twitch** | **roll** |
| 0 | 0% | 0% | 100% | 0% | 7% | 0% | 0% | 0% | 0% | 0% |
| 2 | 0% | 0% | 100% | 0% | 10% | 0% | 0% | 0% | 0% | 0% |
| 4 | 0% | 0% | 100% | 0% | 20% | 0% | 0% | 0% | 0% | 0% |
| 6 | 0% | 0% | 97% | 0% | 0% | 0% | 0% | 0% | 0% | 0% |
| 8 | 50% | 0% | 97% | 0% | 3% | 0% | 0% | 0% | 0% | 0% |
| 10 | 77% | 7% | 90% | 0% | 0% | 0% | 0% | 0% | 0% | 0% |
| 12 | 93% | 77% | 7% | 0% | 0% | 0% | 0% | 0% | 0% | 0% |
| 14 | 87% | 27% | 0% | 0% | 0% | 0% | 0% | 0% | 0% | 0% |
| 16 | 100% | 0% | 0% | 0% | 0% | 0% | 0% | 0% | 0% | 0% |

**Table S5.** *Drosophila pseudoobscura* cold-evoked behavioral program. Each value shows the % of larvae performing a particular behavior, at a particular temperature. See **Figure 1**.

| *willistoni* | | | | | | | | | | |
|---|---|---|---|---|---|---|---|---|---|---|
| **Temp** | **Loco** | **P** | **CT** | **SER** | **HR** | **TR** | **HTR** | **HW** | **twitch** | **roll** |
| 0 | 0% | 0% | 97% | 0% | 0% | 3% | 0% | 0% | 0% | 0% |
| 2 | 0% | 0% | 90% | 0% | 0% | 0% | 0% | 0% | 0% | 3% |
| 4 | 0% | 0% | 90% | 0% | 0% | 7% | 0% | 0% | 0% | 0% |
| 6 | 0% | 0% | 93% | 0% | 0% | 3% | 0% | 0% | 0% | 0% |
| 8 | 0% | 0% | 100% | 0% | 0% | 0% | 0% | 0% | 0% | 0% |
| 10 | 0% | 0% | 90% | 0% | 3% | 0% | 0% | 0% | 0% | 0% |
| 12 | 87% | 87% | 0% | 0% | 0% | 0% | 0% | 0% | 0% | 0% |
| 14 | 100% | 100% | 0% | 0% | 0% | 0% | 0% | 0% | 0% | 0% |
| 16 | 100% | 50% | 0% | 0% | 0% | 0% | 0% | 0% | 0% | 0% |

**Table S6.** *Drosophila willistoni* cold-evoked behavioral program. Each value shows the % of larvae performing a particular behavior, at a particular temperature. See **Figure 1**.

| *equinoxialis* | | | | | | | | | | |
|---|---|---|---|---|---|---|---|---|---|---|
| **Temp** | **Loco** | **P** | **CT** | **SER** | **HR** | **TR** | **HTR** | **HW** | **twitch** | **roll** |
| 0 | 0% | 0% | 100% | 0% | 0% | 10% | 3% | 0% | 0% | 0% |
| 2 | 0% | 0% | 100% | 0% | 0% | 17% | 13% | 0% | 0% | 0% |
| 4 | 0% | 0% | 100% | 0% | 0% | 0% | 0% | 0% | 0% | 0% |
| 6 | 0% | 0% | 73% | 0% | 7% | 3% | 0% | 7% | 3% | 3% |
| 8 | 0% | 0% | 80% | 0% | 10% | 3% | 3% | 10% | 3% | 3% |
| 10 | 10% | 14% | 48% | 0% | 0% | 7% | 3% | 31% | 0% | 0% |
| 12 | 87% | 40% | 20% | 0% | 0% | 3% | 3% | 27% | 0% | 7% |
| 14 | 100% | 67% | 0% | 0% | 0% | 0% | 0% | 0% | 7% | 0% |
| 16 | 100% | 80% | 0% | 0% | 0% | 0% | 0% | 0% | 0% | 0% |

**Table S7.** *Drosophila equinoxialis* cold-evoked behavioral program. Each value shows the % of larvae performing a particular behavior, at a particular temperature. See **Figure 1**.

| *funebris* | | | | | | | | | | |
|---|---|---|---|---|---|---|---|---|---|---|
| **Temp** | **Loco** | **P** | **CT** | **SER** | **HR** | **TR** | **HTR** | **HW** | **twitch** | **roll** |
| 0 | 0% | 0% | 100% | 0% | 0% | 0% | 0% | 0% | 0% | 0% |
| 2 | 0% | 0% | 100% | 0% | 7% | 0% | 0% | 0% | 0% | 0% |
| 4 | 0% | 0% | 83% | 0% | 3% | 0% | 0% | 10% | 0% | 0% |
| 6 | 13% | 0% | 97% | 0% | 0% | 0% | 0% | 3% | 0% | 0% |
| 8 | 13% | 0% | 97% | 0% | 0% | 0% | 0% | 0% | 0% | 0% |
| 10 | 90% | 14% | 17% | 0% | 0% | 0% | 0% | 24% | 0% | 0% |
| 12 | 100% | 0% | 10% | 0% | 3% | 0% | 0% | 20% | 0% | 0% |
| 14 | 100% | 3% | 3% | 0% | 0% | 0% | 0% | 0% | 0% | 0% |
| 16 | 100% | 0% | 0% | 0% | 0% | 0% | 0% | 0% | 0% | 0% |

**Table S8.** *Drosophila funebris* cold-evoked behavioral program. Each value shows the % of larvae performing a particular behavior, at a particular temperature. See **Figure 1**.

| *arizonae* | | | | | | | | | | |
|---|---|---|---|---|---|---|---|---|---|---|
| **Temp** | **Loco** | **P** | **CT** | **SER** | **HR** | **TR** | **HTR** | **HW** | **twitch** | **roll** |
| 0 | 0% | 0% | 100% | 0% | 0% | 10% | 3% | 0% | 0% | 0% |
| 2 | 0% | 0% | 100% | 10% | 0% | 3% | 3% | 0% | 0% | 0% |
| 4 | 0% | 0% | 100% | 23% | 0% | 20% | 13% | 0% | 0% | 0% |
| 6 | 0% | 0% | 100% | 87% | 10% | 30% | 0% | 0% | 3% | 0% |
| 8 | 27% | 0% | 93% | 10% | 3% | 3% | 0% | 3% | 0% | 7% |
| 10 | 53% | 30% | 67% | 0% | 0% | 0% | 0% | 3% | 0% | 0% |
| 12 | 60% | 90% | 0% | 0% | 0% | 0% | 0% | 10% | 0% | 0% |
| 14 | 93% | 33% | 0% | 0% | 0% | 0% | 0% | 17% | 0% | 0% |
| 16 | 100% | 3% | 0% | 0% | 0% | 0% | 0% | 0% | 0% | 0% |

**Table S9.** *Drosophila arizonae* cold-evoked behavioral program. Each value shows the % of larvae performing a particular behavior, at a particular temperature. See **Figure 1**.

***littoralis***

| Temp | Loco | P | CT | SER | HR | TR | HTR | HW | twitch | roll |
|---|---|---|---|---|---|---|---|---|---|---|
| 0 | 0% | 3% | 100% | 0% | 10% | 0% | 0% | 0% | 0% | 0% |
| 2 | 0% | 17% | 100% | 0% | 10% | 3% | 0% | 0% | 0% | 0% |
| 4 | 0% | 0% | 100% | 0% | 10% | 0% | 0% | 0% | 0% | 0% |
| 6 | 13% | 0% | 100% | 0% | 3% | 10% | 0% | 0% | 0% | 0% |
| 8 | 20% | 3% | 93% | 0% | 0% | 0% | 0% | 0% | 0% | 0% |
| 10 | 37% | 7% | 90% | 0% | 3% | 3% | 0% | 0% | 0% | 0% |
| 12 | 63% | 30% | 50% | 0% | 0% | 0% | 0% | 0% | 0% | 0% |
| 14 | 87% | 57% | 13% | 0% | 0% | 3% | 0% | 0% | 0% | 0% |
| 16 | 100% | 20% | 0% | 0% | 3% | 0% | 0% | 0% | 0% | 0% |

**Table S10.** *Drosophila littoralis* cold-evoked behavioral program. Each value shows the % of larvae performing a particular behavior, at a particular temperature.  See **Figure 1**.

| | | | | | *mojavensis* | | | | | |
|---|---|---|---|---|---|---|---|---|---|---|
| **Temp** | **Loco** | **P** | **CT** | **SER** | **HR** | **TR** | **HTR** | **HW** | **twitch** | **roll** |
| 0 | 0% | 0% | 100% | 30% | 3% | 27% | 10% | 0% | 0% | 0% |
| 2 | 0% | 0% | 93% | 17% | 0% | 23% | 7% | 0% | 0% | 0% |
| 4 | 0% | 0% | 97% | 3% | 0% | 20% | 0% | 0% | 0% | 0% |
| 6 | 3% | 3% | 97% | 40% | 3% | 67% | 0% | 0% | 0% | 0% |
| 8 | 37% | 3% | 80% | 13% | 0% | 0% | 0% | 10% | 0% | 0% |
| 10 | 70% | 13% | 30% | 3% | 10% | 0% | 0% | 40% | 0% | 3% |
| 12 | 93% | 0% | 0% | 0% | 3% | 0% | 0% | 0% | 0% | 0% |
| 14 | 100% | 0% | 0% | 0% | 3% | 0% | 0% | 0% | 0% | 3% |
| 16 | 100% | 0% | 0% | 0% | 0% | 0% | 0% | 0% | 0% | 0% |

**Table S11.** *Drosophila mojavensis* cold-evoked behavioral program. Each value shows the % of larvae performing a particular behavior, at a particular temperature. See **Figure 1**.

| *hydei* | | | | | | | | | | |
|---|---|---|---|---|---|---|---|---|---|---|
| **Temp** | **Loco** | **P** | **CT** | **SER** | **HR** | **TR** | **HTR** | **HW** | **twitch** | **roll** |
| 0 | 0% | 0% | 100% | 70% | 0% | 0% | 0% | 0% | 0% | 0% |
| 2 | 0% | 0% | 100% | 33% | 0% | 0% | 0% | 0% | 0% | 0% |
| 4 | 0% | 0% | 100% | 43% | 0% | 0% | 0% | 0% | 0% | 0% |
| 6 | 0% | 0% | 100% | 30% | 0% | 17% | 0% | 0% | 0% | 0% |
| 8 | 3% | 0% | 97% | 0% | 0% | 0% | 0% | 3% | 0% | 0% |
| 10 | 37% | 7% | 53% | 30% | 0% | 0% | 0% | 20% | 0% | 0% |
| 12 | 77% | 87% | 0% | 3% | 0% | 0% | 0% | 10% | 0% | 0% |
| 14 | 93% | 10% | 0% | 0% | 0% | 0% | 0% | 0% | 0% | 0% |
| 16 | 100% | 0% | 0% | 0% | 0% | 0% | 0% | 0% | 0% | 0% |

**Table S12.** *Drosophila hydei* cold-evoked behavioral program. Each value shows the % of larvae performing a particular behavior, at a particular temperature.  See **Figure 1**.
